# Population Genetics of Native Red Mulberry at Its Northwestern Boundary Suggests Postglacial Founder Effects

**DOI:** 10.64898/2026.07.11.737963

**Authors:** Spencer J. Schreier, Madhav P. Nepal

## Abstract

*Morus rubra* is native to the eastern United States, with its range extending into the Upper Midwest and southern Ontario, Canada. Its present distribution suggests that past glacial events in North America may have influenced the genetic structure of populations at the species’ northwestern range boundary. This study assessed genetic variation among six *M. rubra* populations believed to have experienced postglacial colonization using published nuclear microsatellite markers and sequences from the chloroplast *trnL–trnF* region. Five nuclear microsatellite markers previously developed for *M. alba* were successfully transferred to *M. rubra*, while the chloroplast *trnL–trnF* region provided an additional marker for evaluating chlorotype diversity. Nuclear microsatellite diversity was higher in southern unglaciated populations than in northern glaciated populations, a pattern consistent with the observed distribution of chlorotype diversity. Together, these results support ancient founder effects associated with leading-edge expansion following glacial recession and suggest that postglacial colonization contributed to the present-day genetic structure of *M. rubra* at its northwestern range boundary. Because *M. rubra* hybridizes with the naturalized invasive *M. alba*, reduced genetic diversity in marginal populations may increase their vulnerability to genetic swamping. The markers characterized in this study provide useful tools for population genetic research in *Morus*, and the findings have important implications for the conservation and management of marginal and threatened *M. rubra* populations in the Upper Midwest.

## 1. Introduction

*Morus rubra* L. (Moraceae), commonly known as Red Mulberry, is one of the 13 recognized species in the genus *Morus* and is native to the eastern United States, with its distribution extending into the Upper Midwest and southern Ontario, Canada (Nepal & Purintun, 2021) (Nepal, 2008) (Nepal & Ferguson, 2012). As an ecologically important native tree, *M. rubra* supports wildlife, has a long history of nutritional and traditional medicinal use by Native Americans (Moerman, 1986), and has become an increasing conservation concern because of habitat fragmentation and hybridization with the introduced *M. alba* (Burgess et al., 2005).

Evidence from numerous plant species suggests that glacial cycles during the Pleistocene profoundly influenced population structure throughout the Northern Hemisphere (Broyles, 1998; Gamache et al., 2003; Gugerli et al., 2001; Palmé, 2002; Skrede et al., 2006)(Pielou). The present distribution of *M. rubra* has therefore raised the question of whether the Wisconsin glaciation and subsequent recession of the Laurentide Ice Sheet shaped the genetic structure of populations at the species’ northwestern range boundary. During the last glacial maximum, the Laurentide Ice Sheet extended to approximately 40–50°N across North America (Ingólfsson, 2008), eliminating vegetation over large areas. Analysis of the oxygen isotopes in deep sea cores indicated that such glacial events followed a cyclic pattern: long glaciation phases were followed by drastic increases in temperature that caused short periods of glacial recession (Broecker & Donk, 1970; Davis, 1983). Such recession provided new niches for surviving populations of *M. rubra* to colonize. This colonization may have shaped the genetic structure of the current *M. rubra* populations in the Upper Midwest. Evidence suggest that when biotic or abiotic factors influence the movement of individuals into open niches far away from their source population, allelic frequencies may fluctuate across the expanded population range (Gamache et al., 2003). In a glacier leading-edge expansion, the *M. rubra* individuals from the leading-edge subpopulations would not carry all the alleles from the source population (Bergstrom, 2012; Raven, 2010), resulting in less genetically-diverse populations and a selective effect on genetic variation (Broyles, 1998; Hewitt, 1996). This colonization is referred to as leading edge expansion, a mechanism of founder effect that results in new populations with tapered genetic diversity as expected of the initial leading-edge colonizers (Bergstrom, 2012). Similar glaciation induced founder effect may have had a critical influence on the current genetic structure of *M. rubra* populations. Examination of DNA evidence can determine the overall contribution of the glacier-induced leading-edge expansion and more recent gene flow to current *M. rubra* populations.

The evolutionary history and population structure of a species can be investigated using nuclear and chloroplast DNA markers (Hewitt, 2004). Because chloroplast DNA is maternally inherited, highly conserved, and evolves slowly (Hartl & Clark, 2007) (Wolfe et al., 1987), it is particularly useful for reconstructing historical migration routes and postglacial recolonization (Dumolin et al., 1995; Gamache et al., 2003; Heuertz et al., 2004). In contrast, biparentally inherited nuclear markers provide greater resolution of contemporary population structure and gene flow (McPherson et al., 2013). Together, chloroplast and nuclear markers provide complementary perspectives on historical colonization and present-day genetic diversity. Applying these complementary molecular markers to *M. rubra* provides an opportunity to reconstruct its postglacial history while evaluating the genetic consequences of range expansion in a species of conservation concern (Burgess et al., 2008). *Morus rubra* is currently considered endangered in Canada (Burgess et al., 2008) and in Connecticut and Massachusetts, and threatened in Michigan and Vermont in the United States (Sullivan, 1993). In addition, *M. rubra* hybridizes readily with the introduced invasive *M. alba* (Burgess et al., 2005), introduced in North America from Asia during the colonial era (Stone, 2009), increasing the risk of genetic swamping in small or isolated populations. We hypothesized that repeated recession of the Laurentide Ice Sheet generated a cascade of leading-edge colonization events, resulting in younger populations with reduced genetic diversity toward the northwestern range of *M. rubra*. To test this hypothesis, we evaluated the interspecific transferability of seven nuclear microsatellite markers and characterized the nuclear and chloroplast genetic diversity of six M. rubra populations in the Upper Midwest.

## 2. Materials and Methods

### 2.1. DNA Extraction and PCR Screening

Fresh leaf samples were collected from 10–18 randomly selected *Morus rubra* individuals in each of six populations from Kansas, Iowa, Wisconsin, and Nebraska (Table 1). Leaves were dried in silica gel, and genomic DNA was extracted using the DNeasy Plant Mini Kit (Qiagen Corp., Valencia, CA) following the manufacturer’s protocol. A total of 12 nuclear microsatellite markers previously developed for *M. indica* (= *M. alba*) and *M. boninensis* were screened for cross-species amplification and transferability to *M. rubra* (Aggarwal et al., 2004) (Tani et al., 2005). PCR conditions were optimized separately for each marker set. The MulSTR markers were amplified in 15-μL reactions containing 25 ng genomic DNA, 3 μL of 5× GoTaq Flexi buffer, 1.5 μL of 2.5 mM MgCl_2_, 1.2 μL of 10 mM dNTP mixture, 0.38 μL each of 10 pM forward and reverse primers, and 1 U of GoTaq DNA polymerase. Amplification consisted of an initial denaturation at 94°C for 2 min; 30 cycles of 94°C for 45 s, 51°C for 1 min, and 72°C for 2 min; and a final extension at 72°C for 5 min. Five nuclear microsatellite markers produced clear and reproducible amplification and were retained for genotyping and population genetic analyses, whereas the remaining markers did not amplify consistently in *M. rubra*. The chloroplast *trnL–trnF* intergenic spacer was amplified following Nepal and Ferguson (2012).

**Table 1.**
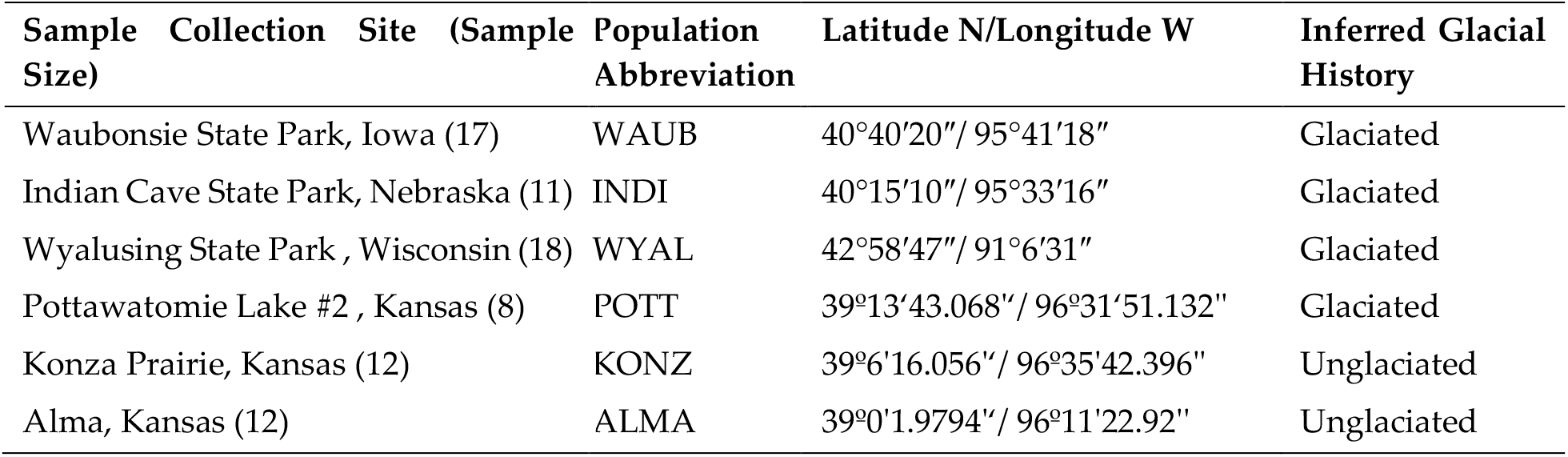
Sampling locations, population abbreviations, geographic coordinates, and inferred glacial histories of six *Morus rubra* populations from the Upper Midwest United States. Populations were classified as originating from either glaciated or unglaciated regions based on their location relative to the southern extent of Pleistocene glaciation. Sample sizes for each population are given in parentheses following the collection site name. Latitude and longitude coordinates are provided for each sampling location.

PCR was performed in 15-μL reactions containing 25 ng genomic DNA, 5 μL of 5× buffer, 1 μL of dNTPs, 2.5 μL of 2.5 mM MgCl_2_, 2.5 μL of 10 pM primers, and 2 U of GoTaq DNA polymerase. Cycling conditions consisted of an initial denaturation at 94°C for 5 min; 27 cycles of 94°C for 1 min, 55°C for 1 min, and 72°C for 2 min; and a final extension at 72°C for 10 min. Nuclear and chloroplast PCR products were verified by electrophoresis on 1.2% agarose gels stained with ethidium bromide and visualized under ultraviolet light. Thus, the final dataset included six molecular markers comprising five nuclear microsatellite loci and the chloroplast *trnL–trnF* region (Table 2).

**Table 2.**
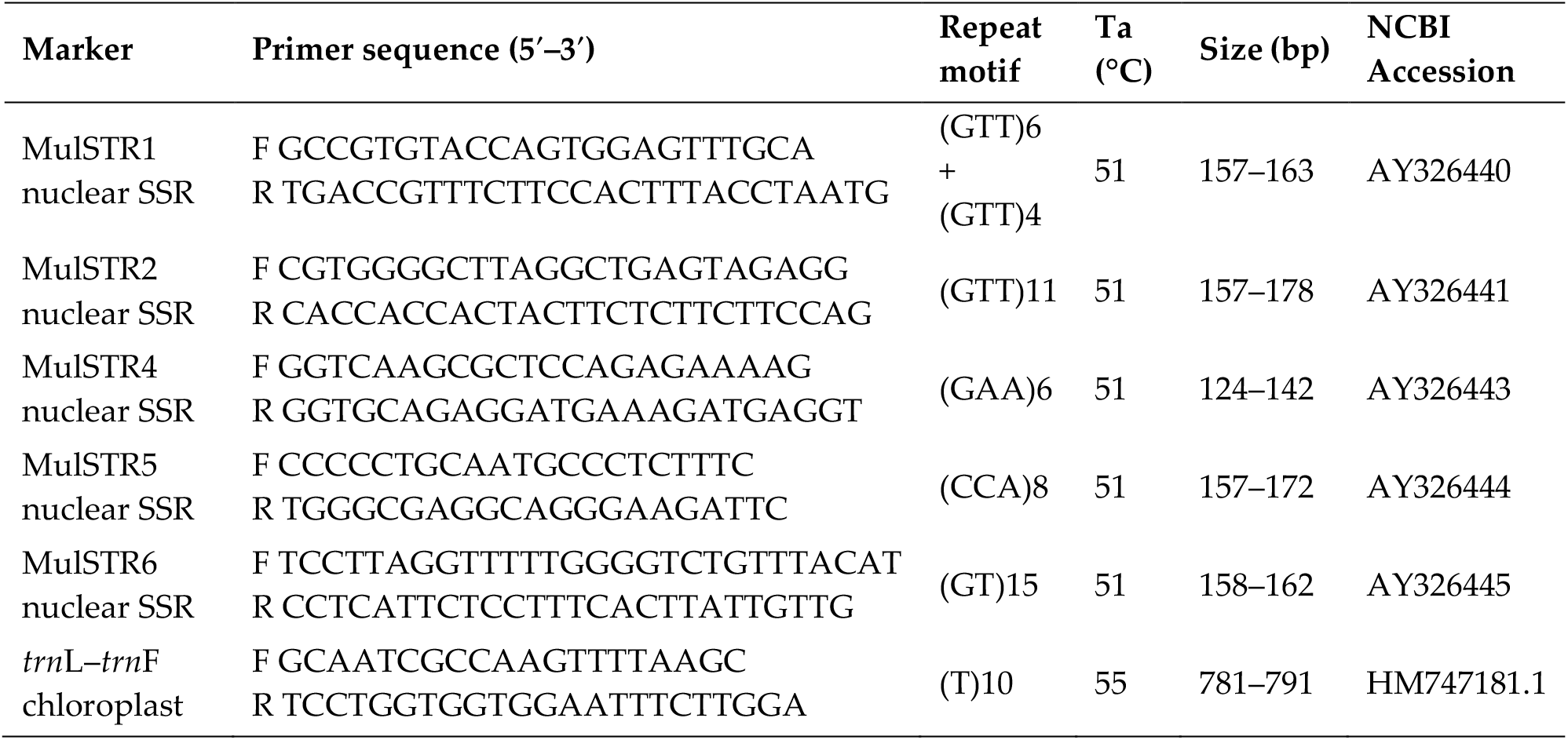
Characteristics of five nuclear microsatellite loci and the chloroplast *trn*L–*trn*F region retained for genetic analyses of *Morus rubra*. Primer sequences, modified from Aggarwal (2004), are shown in the 5′–3′ direction. Ta = annealing temperature; bp = base pairs; SSR = simple sequence repeat.

### 2.2. Microsatellite Genotyping

The PCR conditions described above were used for microsatellite genotyping, with an M13-tailed forward primer and FAM-labeled universal primer incorporated for fluorescent fragment analysis. Genotyping was performed at the Iowa State University DNA Facility using an ABI 3730xl DNA Analyzer (Applied Biosystems). A total of 78 *M. rubra* individuals were successfully genotyped at the five transferable MulSTR loci. Allele sizes were scored and allele reports generated using GeneMarker v2.4.0 (SoftGenetics).

### 2.3. Chloroplast Sequencing

The amplified trnL–trnF region was purified using the QIAquick PCR Purification Kit (Qiagen Corp., Valencia, CA) and sequenced at the Iowa State University DNA Facility. Sequences from 87 individuals were edited in Sequencher v5.0 (Gene Codes Corporation) and aligned in ClustalX (Thompson et al., 2001) to identify sequence polymorphisms and define chlorotypes.

### 2.4. Genetic Diversity

Results from GeneMarker V2.4.0 (Softgenetics) were examined for genetic diversity. Evaluation of linkage disequilibrium and deviation from Hardy-Weinberg Equilibrium (H.W.E.) for all loci was performed in Arlequin V3.1 (Excoffier et al., 2005). Genetic diversity of each locus was estimated by its polymorphism (P), allele count, and heterozygosity using Arlequin V3.1 (Excoffier et al., 2005). Within each population, the microsatellites expressed varied allelic richness (A) and heterozygosity levels (Ho; He). These values were averaged to determine the genetic diversity indices for each population, namely, allele count and heterozygosity. To examine the presence of inbreeding, the inbreeding coefficient (F) for each population determined by averaging the locus’ F values in FSTAT (Goudet, 2002).

### 2.5. Population Genetic Structure

Population structure was analyzed for the segregation of population genetic structures. A genetic distance dendrogram was constructed to facilitate population structure inference. Nei’s genetic diversity indexes (Masatoshi Nei, 1973) were calculated to determine gene differentiation (Gst). Total gene diversity (Ht), gene diversity within populations (Hs), and gene diversity among populations (Dst) were determined using the statistical program FSTAT (Goudet, 2002). An AMOVA was performed using the Fst analysis of Arlequin V3.1 (Excoffier et al., 2005) to identify differentiation within populations and among populations (Fst). Fst and Gst values were later observed together, as these values were expected to exhibit a close correlation. A Bayesian model replica was performed to infer population allele composition and theoretical ancestry based on individual genotypes at multiple loci using STRUCTURE (Pritchard et al., 2000). Allele ancestry and frequency within K=1-9 populations were examined with a burn in period of 10,000 and 50,000 Markov chain Monte Carlo (MCMC) algorithm repetitions after each burn-in. 20 repetitions were completed for each value of K. ΔK was determined to to be 4, based on the method demonstrated by Evanno et al. (Evanno et al., 2005). PopGen32 (Yeh et al., 2000) was used to create a dendrogram of the sampled populations based on Nei’s genetic distance (D) (M. Nei, 1973), and using an UPGMA method to group populations together.

### 2.6. Chlorotype Analysis

Analysis of the chloroplast DNA focused on the number of chlorotypes produced by cpDNA sequence polymorphism. Sequenced cpDNA results were re-examined using SEQUENCHER version 5.0 (GeneCodesCorporation) to edit unclear sequence results. SEQUENCHER editing fidelity was verified by comparison with ClustalX (Larkin et al., 2007) results, prior to analysis of potential chlorotypes among individuals. ClustalX (Larkin et al., 2007) displayed the amplified sequences in parallel and their SNPs. Sites with SNPs were cross-referenced by examining nucleotide indels, transversions, and transitions. With these SNPs identified, a number of chlorotypes were identified and compared to chlorotype diversity by geographic location of each population, to observe correlation between latitude and chlorotype diversity. The data matrix was analyzed using the programs DnaSP (Librado & Rozas, 2009) and Network 4.1.0.9 (www.fluxus-engineering.com). The software DnaSP (Librado & Rozas, 2009) evaluated the haplotype diversity of the amplified chloroplast region and the nucleotide diversity (π). Network 4.1.0.9 was used to construct a mutation-distance map of the chlorotypes, which was then used for evaluating the relationship between the various chlorotypes.

## 3. Results

### 3.1. Marker Transferability and Locus Level Genetic Diversity

Of the 12 nuclear microsatellite markers screened, five MulSTR loci produced clear and reproducible genotypes in *M. rubra* and were subjected to population structure analyses. Some individuals displayed more than two allelic peaks at one or more loci, indicating multiallelic profiles that require cautious interpretation. All five loci were polymorphic across the sampled *M. rubra* populations. The number of alleles per locus ranged from 5 to 6, observed heterozygosity (H_O_) from 0.21 to 0.50, and expected heterozygosity (H_E_) from 0.52 to 0.76 (Table 3). Total gene diversity (H_t_) ranged from 0.65 to 0.84, with a mean of 0.76, whereas mean within-population diversity (H_s_) ranged from 0.54 to 0.78, with an overall mean of 0.66. Gene diversity among populations (D_st_) ranged from 0.06 to 0.12, and G__st_ ranged from 0.07 to 0.17, averaging 0.13. H_O_ was lower than H_E_ at every locus, and each locus deviated significantly from Hardy–Weinberg expectations in at least two populations; MulSTR1 and MulSTR6 deviated in all six populations (Table 3).

**Table 3.**
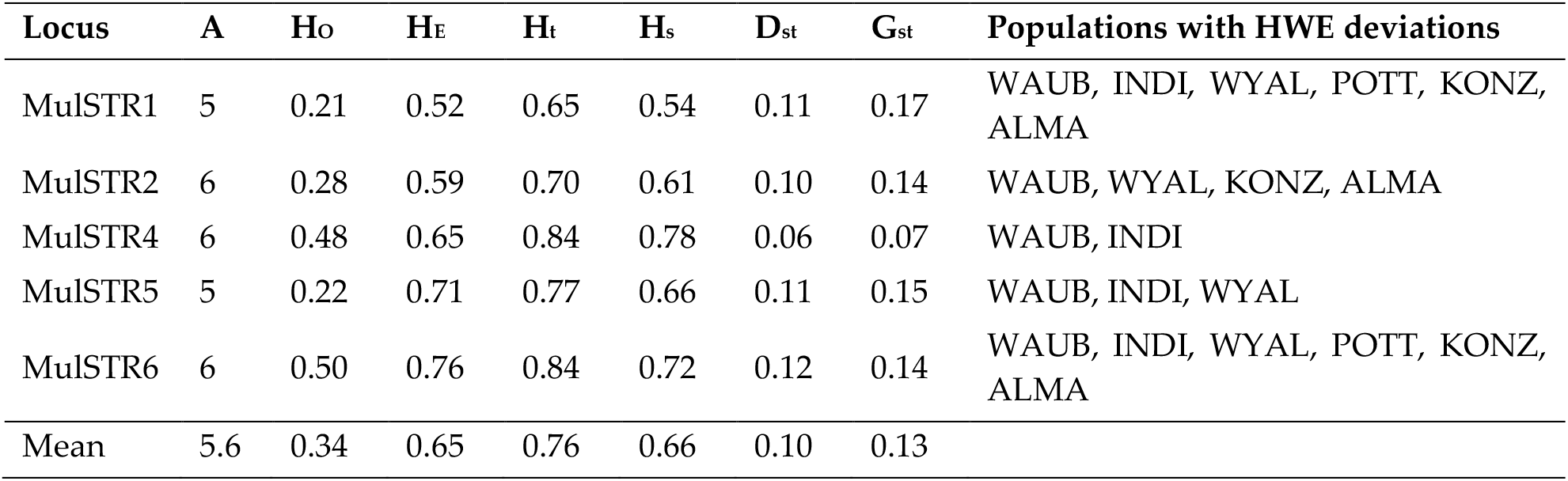
Locus level nuclear microsatellite diversity and population differentiation in *Morus rubra*. A, number of alleles; H_O_, observed heterozygosity; H_E_, expected heterozygosity; H_t_, total gene diversity; H_s_, mean within-population gene diversity; D_st_, gene diversity among populations; G_st_, coefficient of genetic differentiation. The final column lists populations with significant departures from Hardy–Weinberg equilibrium at p < 0.05.

| Locus | A | H<sub>O</sub> | H<sub>E</sub> | H<sub>T</sub> | H<sub>S</sub> | D<sub>st</sub> | G<sub>st</sub> | Populations with HWE deviations |
| --- | --- | --- | --- | --- | --- | --- | --- | --- |
| MulSTR1 | 5 | 0.21 | 0.52 | 0.65 | 0.54 | 0.11 | 0.17 | WAUB, INDI, WYAL, POTT, KONZ, ALMA |
| MulSTR2 | 6 | 0.28 | 0.59 | 0.70 | 0.61 | 0.10 | 0.14 | WAUB, WYAL, KONZ, ALMA |
| MulSTR4 | 6 | 0.48 | 0.65 | 0.84 | 0.78 | 0.06 | 0.07 | WAUB, INDI |
| MulSTR5 | 5 | 0.22 | 0.71 | 0.77 | 0.66 | 0.11 | 0.15 | WAUB, INDI, WYAL |
| MulSTR6 | 6 | 0.50 | 0.76 | 0.84 | 0.72 | 0.12 | 0.14 | WAUB, INDI, WYAL, POTT, KONZ, ALMA |
| Mean | 5.6 | 0.34 | 0.65 | 0.76 | 0.66 | 0.10 | 0.13 |  |

### 3.2. Population Genetic Diversity and Structure

Population-level diversity differed markedly between populations from glaciated and unglaciated regions (Table 4; Figure 1). In the four glaciated populations, the mean number of alleles per locus ranged from 2 to 4, H_O_ from 0.11 to 0.25, H_E_ from 0.47 to 0.58, and chlorotype richness from 2 to 4. In contrast, the two unglaciated populations had 9–10 alleles per locus, H_O_ of 0.57–0.63, H_E_ of 0.89–0.90, and 4–5 chlorotypes. Averaged by glacial history, glaciated populations had lower mean allele number (3.0 vs. 9.5), H_O_ (0.208 vs. 0.600), H_E_ (0.523 vs. 0.895), and chlorotype richness (3.0 vs. 4.5) than unglaciated populations. All populations showed positive F__IS_ values, ranging from 0.307 in ALMA to 0.800 in WAUB; the mean F_IS_ was 0.616 for glaciated populations and 0.341 for unglaciated populations.

**Table 4.** Population level nuclear and chloroplast diversity in six *Morus rubra* populations. Population history indicates whether the collection site was glaciated (G) or unglaciated (U) during the last glacial maximum. N = sample size; A = mean number of nuclear alleles per locus; P = percentage of polymorphic nuclear loci; H_O_ = observed heterozygosity; H_E_ = expected heterozygosity; F_IS_ = within-population inbreeding coefficient; h = chlorotype richness. Chlorotypes were identified from the chloroplast *trn*L–*trn*F region.

| Population history | Sample Size (N) | A | P (%) | H<sub>O</sub> | H<sub>E</sub> | F<sub>IS</sub> | Chlorotypes (h) |
| --- | --- | --- | --- | --- | --- | --- | --- |
| WAUB (G) | 17 | 3 | 80 | 0.11 | 0.52 | 0.800 | CP6, CP7, CP8, CP10 (4) |
| INDI (G) | 11 | 3 | 80 | 0.24 | 0.52 | 0.541 | CP1, CP6, CP7 (3) |
| WYAL (G) | 18 | 4 | 100 | 0.25 | 0.58 | 0.585 | CP5, CP6 (2) |
| POTT (G) | 8 | 2 | 100 | 0.23 | 0.47 | 0.537 | CP2, CP6, CP9 (3) |
| KONZ (U) | 12 | 9 | 100 | 0.57 | 0.89 | 0.374 | CP3, CP4, CP6, CP11, CP12 (5) |
| ALMA (U) | 12 | 10 | 100 | 0.63 | 0.90 | 0.307 | CP3, CP6, CP11, CP12 (4) |

**Figure 1.**
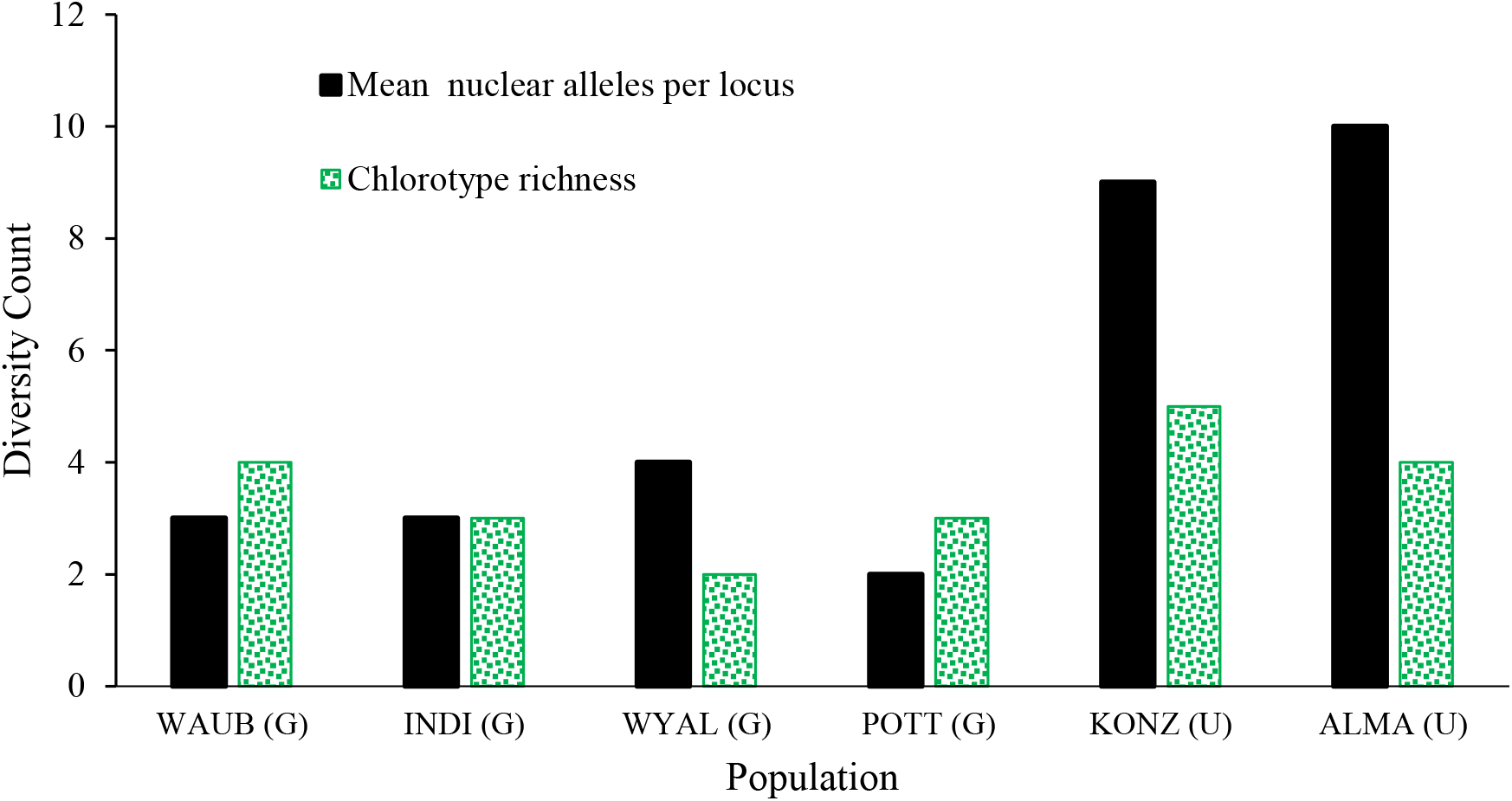
Nuclear allelic diversity and chlorotype richness across six *Morus rubra* populations. Mean allele number per nuclear microsatellite locus and the number of *trn*L–*trn*F chlorotypes are shown for each population. Populations are ordered according to their inferred glacial history. The first four populations on the left are glaciated (G), whereas the remaining populations on the right are unglaciated (U).

AMOVA attributed 83.6% of the nuclear microsatellite variation to differences within populations and 16.4% to differences among populations, producing an overall F_ST_ of 0.1649 (**Table 5**). Bayesian clustering supported K = 4. KONZ and ALMA populations displayed similar ancestry profiles that were distinct from the four glaciated populations. INDI and WYAL also showed broadly similar allele-frequency composition, POTT was dominated by one ancestry component, and WAUB included admixture components also represented in the southern populations (**Figure 2**).

**Table 5.** Analysis of molecular variance for six *Morus rubra* populations based on five nuclear microsatellite loci. d.f. = degrees of freedom; F_ST_ = fixation index measuring differentiation among populations.

| Source of variation | d.f. | Sum of squares | Variance component | Variation (%) | $F_{ST}$ |
| --- | --- | --- | --- | --- | --- |
| Among populations | 5 | 48.31 | 0.31 | 16.4 | 0.1649 |
| Within populations | 150 | 239.15 | 1.59 | 83.6 | — |
| Total | 155 | 287.46 | 1.91 | 100.0 | — |

**Figure 2.**
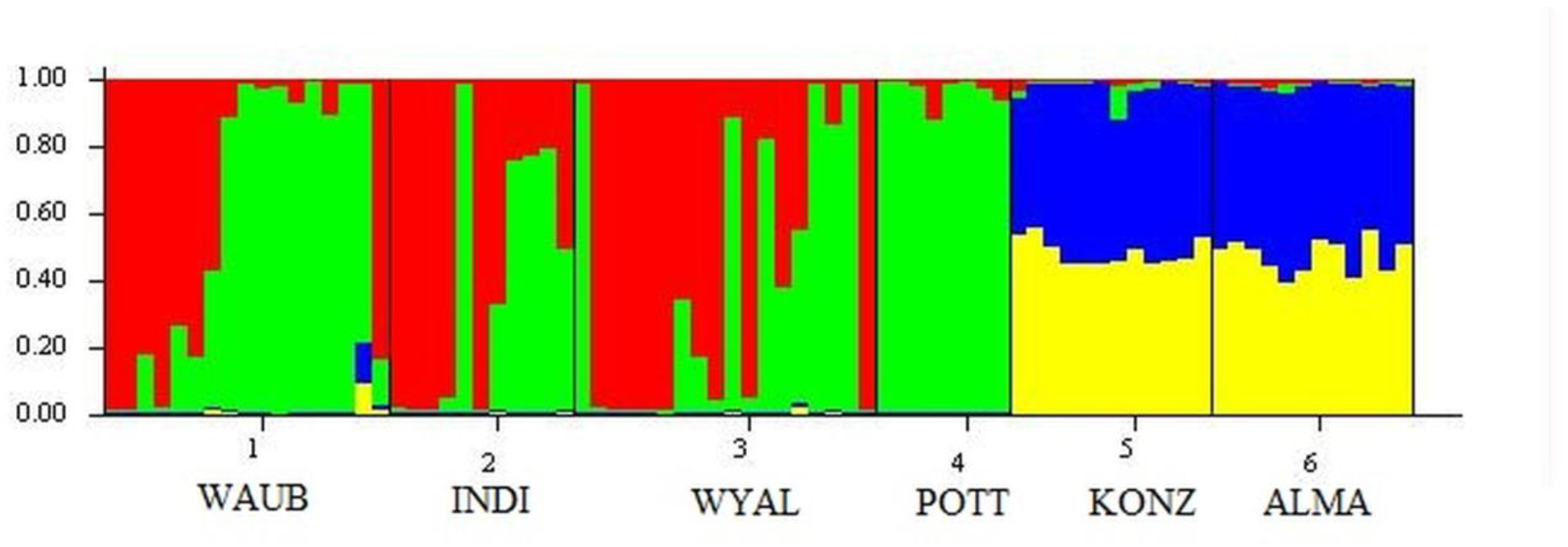
Bayesian population structure of six *Morus rubra* populations at K = 4. Each vertical bar represents an individual, and colored segments represent estimated membership coefficients in four genetic clusters. Individuals are grouped by population in the order WAUB, INDI, WYAL, POTT, KONZ, and ALMA.

The UPGMA dendrogram based on Nei’s genetic distance recovered three population pairs: WAUB & POTT, INDI & WYAL, and KONZ & ALMA. The two unglaciated populations formed a separate major cluster from the four glaciated populations (**Figure 3**).

**Figure 3.**
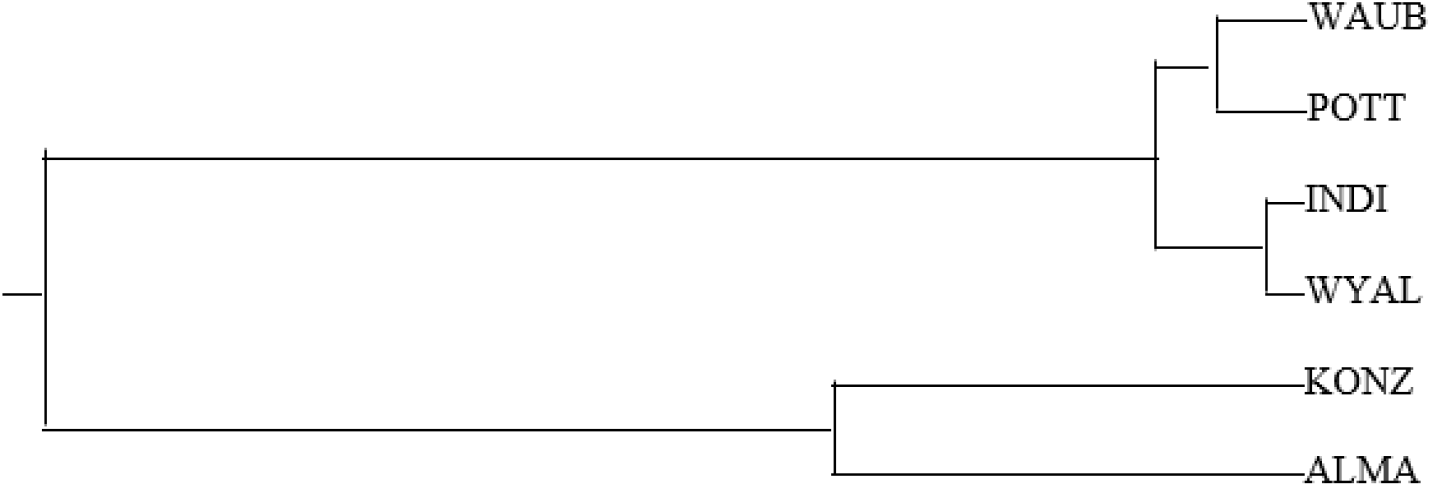
UPGMA dendrogram based on Nei’s genetic distance among six *Morus rubra* populations. The analysis grouped WAUB with POTT, INDI with WYAL, and the two unglaciated populations KONZ and ALMA.

### 3.3. Chloroplast DNA Diversity

The 946-bp *trn*L–*trn*F alignment from 87 individuals contained 12 variable sites that defined 12 chlorotypes. Overall haplotype diversity was 0.33 and nucleotide diversity was 0.00052. Chlorotype richness ranged from two in WYAL to five in KONZ (Table 4), and CP6 occurred in all six populations. The chlorotype network connected the widespread CP6 lineage with multiple lower-frequency chlorotypes separated by one to several mutational steps (**Figure 4**).

**Figure 4.**
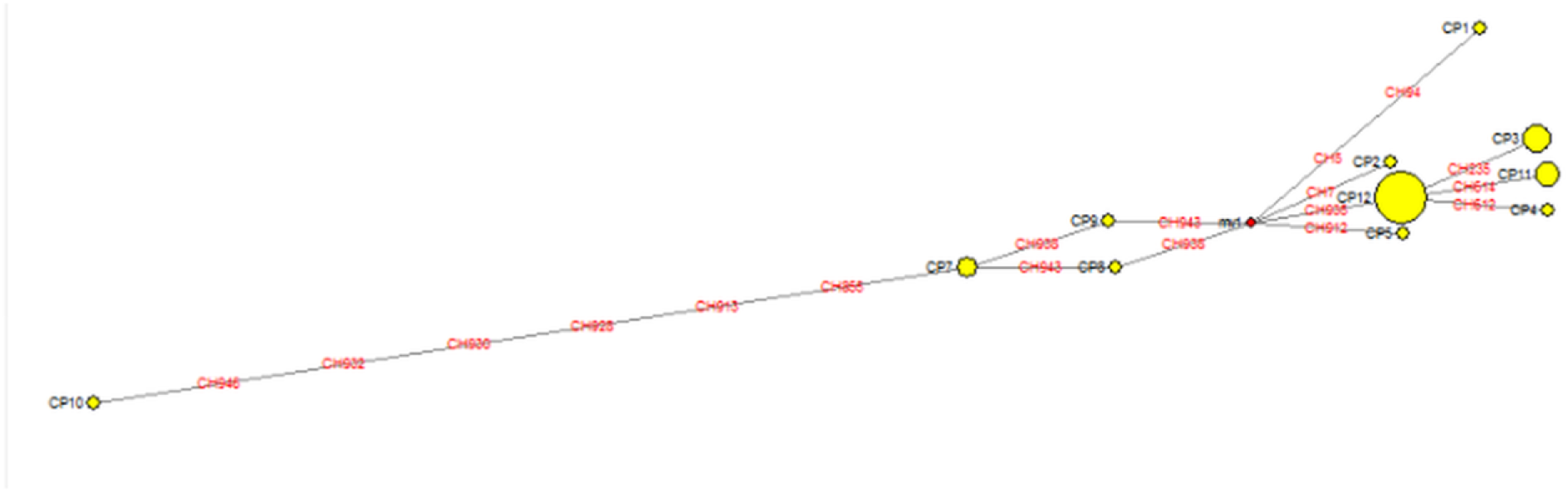
Chlorotype network for 12 *Morus rubra* chlorotypes identified from the chloroplast trnL–trnF region. Node size reflects relative chlorotype frequency, and connecting branches represent mutational differences among chlorotypes. CP6 was detected in all six sampled populations.

## 4. Discussion

The nuclear microsatellite and chloroplast sequence data suggest that historical glaciation and subsequent range expansion may have contributed to the current genetic structure of M. rubra near its northwestern boundary. Both markers showed lower genetic diversity in populations from formerly glaciated regions compared with the two southern unglaciated populations, a pattern consistent with expectations under postglacial colonization and founder effects. However, the absence of a clear south-to-north gradient and the limited number of populations and loci examined constrain the strength of inference. Consequently, these results should be viewed as broadly associated with a postglacial founder-effect scenario rather than as conclusive evidence of the evolutionary processes responsible for the observed genetic patterns.

### 4.1. Microsatellite Transferability and Heterozygote Deficiency

Cross-species amplification of microsatellite loci is a cost-effective alternative to developing species-specific markers and has been widely applied in plant genetic studies (Varshney et al., 2005). Five of the 12 nuclear markers screened here amplified consistently in *M. rubra*, demonstrating the transferability of loci developed for other *Morus* species. Transferability most directly reflects conservation of primer-binding regions and should not, by itself, be interpreted as evidence of low genome-wide differentiation among *Morus* species. The five selected loci nevertheless provided sufficient polymorphism to distinguish population-level differences in diversity and structure.

Observed heterozygosity was lower than expected heterozygosity at every locus and in every population. The H_E values of 0.52–0.76 were also lower than the 0.85–0.90 values reported for these markers in other *Morus* material (Aggarwal et al., 2004). These deficits may reflect inbreeding and population subdivision, but null alleles, allele dropout, and duplicated loci associated with cross-species amplification can produce similar patterns. Likewise, the multiallelic profiles observed in some individuals are compatible with gene duplication or polyploidization (Pazy & Zohary, 1965; Zohary & Feldman, 1962), but they do not independently demonstrate allopolyploidy or introgression from *M. alba*. Recent phylogenomic studies have documented reticulation, incomplete lineage sorting, and cytonuclear discordance across *Morus*, reinforcing the need to evaluate possible hybrids with species-diagnostic nuclear SNPs or genome-scale data (Wang et al., 2024; Yang et al., 2023).

### 4.2. Evidence for Postglacial Founder Effects

The strongest result was the concordant reduction of nuclear and chloroplast diversity in the glaciated populations. Mean nuclear allele number was 3.0 in glaciated populations compared with 9.5 in unglaciated populations, mean H_E was 0.523 compared with 0.895, and mean chlorotype richness was 3.0 compared with 4.5. This pattern is consistent with leading-edge expansion, in which repeated colonization by subsets of a source population reduces allelic and haplotypic diversity toward the range margin (Broyles, 1998; Gamache et al., 2003; Gugerli et al., 2001; Hewitt, 1996; Palmé, 2002). A recent synthesis of nuclear microsatellite studies in European beech similarly recovered decreasing genetic diversity with increasing distance from glacial refugia, illustrating the persistence of postglacial demographic signatures in long-lived forest trees (Stefanini et al., 2023)

The STRUCTURE and UPGMA analyses also separated the two unglaciated Kansas populations from the glaciated populations, while AMOVA detected moderate among-population differentiation (FST = 0.1649) despite most variation occurring within populations (83.6%). The high diversity and close clustering of KONZ and ALMA are therefore consistent with these southern populations representing reservoirs of diversity from which northward colonization could have proceeded. However, the present sampling does not establish that they were the direct historical sources of WAUB, INDI, WYAL, and POTT. Range-edge demography, current habitat fragmentation, and historical recolonization can generate similar spatial patterns; a broad analysis of North American plants emphasized that both glacial history and central-marginal demographic processes should be considered when interpreting reduced diversity at northern margins (López-Delgado & Meirmans, 2022).

### 4.3. Concordance and Differences Between Nuclear and Chloroplast Patterns

The nuclear and chloroplast markers were broadly concordant in distinguishing the more diverse unglaciated populations from the glaciated populations. The UPGMA groupings agreed with major similarities in the STRUCTURE analysis, yet nuclear heterozygosity did not always decline with latitude. Similarly, the glaciated WAUB population contained four chlorotypes, equal to the richness observed in unglaciated ALMA. These departures may reflect contemporary gene flow, recombination in the nuclear genome, lateral colonization from unsampled populations, or disturbance-related movement of propagules.

Maternal introgression from *M. alba* could also alter the geographic distribution of chlorotypes because hybrid fitness depends partly on the direction of the cross (Burgess & Husband, 2004). Anthropogenic disturbance has been shown to complicate simple relationships between geography and genetic diversity in recently colonized landscapes (Müller et al., 2012). Thus, the WAUB pattern and the lack of strict latitudinal decline should not be viewed as contradictions of the founder-effect hypothesis, but as evidence that postglacial history has subsequently been modified by gene flow, hybridization, and local demographic processes. Although nucleotide diversity in the *trn*L–*trn*F region was low (π = 0.00052), small differences in chloroplast sequences can retain useful historical information because plant chloroplast DNA generally evolves slowly and is highly conserved (Curtis & Clegg, 1984; Palmer, 1987; Wolfe et al., 1987). The occurrence of CP6 in every population and its connection to multiple chlorotypes in the network are consistent with a widespread ancestral or early colonizing lineage, but the data do not establish CP6 as the sole primary colonizer. A recent whole-plastome analysis of *M. rubra* across a broader geographic sample also recovered 12 haplotypes and identified multiple variable intergenic regions suitable for marker development (Adhikari et al., 2025). That genomic evidence supports the value of chloroplast variation while indicating that whole-plastome or multilocus chloroplast data will provide greater resolution than *trn*L–*trn*F alone.

### 4.4. Conservation Implications and Study Limitations

Positive F_IS_ values occurred in all populations, and WAUB combined the highest F_IS_ (0.800) with the lowest H_O_ (0.11). Because heterozygote deficits can also arise from null alleles or population subdivision, these values should not be interpreted as direct estimates of mating among close relatives. Nevertheless, the combination of low allelic diversity, high heterozygote deficiency, and geographic marginality identifies WAUB and other glaciated populations as priorities for further genetic assessment. Conservation should retain multiple populations rather than focusing only on the most diverse southern sites, because northern populations may contain distinct chlorotypes and genetic combinations produced by postglacial history.

These concerns are heightened by hybridization with the more abundant invasive *M. alba*. Interspecific hybridization can reduce the persistence of rare species (Levin et al., 1996), accelerate displacement by invasive congeners (Huxel, 1999), and alternative population size and genetic composition through recurrent gene flow(Haygood et al., 2003). When fertile hybrids backcross predominantly with the rare species, genetic swamping and assimilation may follow (Ellstrand, 1992; Rieseberg et al., 1989). This risk is well documented for *M. rubra* and *M. alba* (Burgess et al., 2005), and the ecological loss of *M. rubra* may affect associated native forest species (Ambrose & Kirk, 2004). Management should therefore combine protection of genetically diverse populations, preservation of distinct northern lineages, verification of species identity, and monitoring of hybridization.

The present study was limited to six populations, five transferable nuclear loci, uneven sample sizes caused by the rare and patchy distribution of *M. rubra*, and the absence of reference *M. alba* and confirmed hybrid genotypes. Additional highly informative nuclear markers are therefore needed to resolve the extent, directionality, and demographic consequences of introgression. Recent work from our laboratory has generated complete chloroplast (Adhikari et al., 2025) and mitochondrial (Adhikari et al., 2026) genomes, from which hundreds of candidate organellar markers are undergoing validation. Low-coverage Illumina genome-skimming data are also available, and development of a high-quality nuclear reference genome is underway using PacBio long-read and Hi-C scaffolding data. Integrating these resources with expanded range-wide sampling of *M. rubra, M. alba*, and putative hybrids will improve estimates of gene flow, identify distinct conservation units, and strengthen strategies for the restoration and long-term management of native *M. rubra* populations.

## 5. Conclusions

Nuclear microsatellite genotype and chloroplast sequence variation showed concordant reductions in genetic diversity in populations from formerly glaciated regions compared with the two southern unglaciated populations. Glaciated populations averaged 3.0 nuclear alleles per locus, H_O_ of 0.208, H_E_ of 0.523, and three chlorotypes, whereas unglaciated populations averaged 9.5 alleles, H_O_ of 0.600, H_E_ of 0.895, and 4.5 chlorotypes. Together with the STRUCTURE, UPGMA, and AMOVA results, these patterns are consistent with founder effects during postglacial leading-edge expansion, although broader sampling and demographic analyses are needed to distinguish historical colonization from contemporary range-edge processes. All populations showed heterozygote deficiencies, with the highest F_IS observed in WAUB. These marginal populations may be particularly vulnerable to habitat fragmentation and genetic swamping through hybridization with invasive *M. alba*. Conservation efforts should preserve multiple populations, protect genetically diverse southern populations as potential reservoirs, retain distinct northern lineages, and use species-diagnostic markers to identify pure *M. rubra* and hybrids. The five transferable nuclear microsatellite loci and chloroplast *trn*L–*trn*F region provide an initial toolkit for these efforts, while genomic markers will offer greater resolution for future conservation and phylogeographic studies.

## Acknowledgments

S. J. Schreier received the Excellence in Research Award from Van D. and Barbara B. Fishback Honors College, South Dakota State University. Funding for this research was provided by the South Dakota Agricultural Experiment Station Hatch Project (Award No. SD00H800-23) to M.P. Nepal and by an Honors College Undergraduate Research Award to S. J. Schreier. This work was conducted as part of Schreier’s participation in BIOL 498 (Undergraduate Research and Scholarship) under the mentorship of M. P. Nepal in the SDSTATE Department of Biology and Microbiology.

## Author Contributions

MPN conceived and designed the experiments; SJS performed the experiments; MPN and SJS analyzed the data; MN and SJS wrote and revised the manuscript.

## Availability of Data and Materials

The authors confirm that the data supporting the findings of this study are available within the article.

## Ethics Approval

Not applicable

